# Multi-modal Nonlinear Optical and Thermal Imaging Platform for Label-Free Characterization of Biological Tissue

**DOI:** 10.1101/2020.04.06.023820

**Authors:** Wilson R Adams, Brian Mehl, Eric Lieser, Manqing Wang, Shane Patton, Graham A Throckmorton, J Logan Jenkins, Jeremy B Ford, Rekha Gautam, Jeff Brooker, E. Duco Jansen, Anita Mahadevan-Jansen

**Affiliations:** Vanderbilt University, Dept of Biomedical Engineering, Nashville, TN, 37235, USA; Thorlabs Imaging Research, Sterling, VA, USA; Chongqing University, College of Bioengineering, Chongqing, China; Vanderbilt University Medical Center, Dept of Neurosurgery, Nashville, TN, 37232, USA

## Abstract

The ability to characterize the combined structural, functional, and thermal properties of biophysically dynamic samples is needed to address critical questions related to tissue structure, physiological dynamics, and disease progression. Towards this, we have developed an imaging platform that enables multiple nonlinear imaging modalities to be combined with thermal imaging on a common sample. Here we demonstrate label-free multimodal imaging of live cells, excised tissues, and live rodent brain models. While potential applications of this technology are wide-ranging, we expect it to be especially useful in addressing biomedical research questions aimed at the biomolecular and biophysical properties of tissue and their physiology.

## Main Text

With increased focus on high-speed biological processes at molecular, chemical, and biophysical levels, there has come a critical need to develop tools that can keep pace with the number of active research areas in biomedicine. Technical developments in nonlinear optical microscopy have revolutionized our ability to study biophysical and biochemical properties of tissues, both in terms of the wide range of samples, improved contrast modalities, and unprecedented speeds at which imaging is now possible. However, combining direct spatial temperature measurements alongside multiphoton modes of contrast have not been widely adopted. Multiphoton imaging approaches have enabled deep-tissue imaging across numerous model systems, gleaning important insights on the structure and function of cells and tissues (*1*, *2*) while thermal imaging can yield complementary information to relate to study metabolism, circulation, and immune response. Temperature plays a crucial role in biology yet has yet to be extensively demonstrated in any substantial capacity towards biological microscopy applications. Thus, instrumentation to explore the structural, functional, and thermal properties of tissue would prove useful in studying biophysical dynamics in tissue physiology.

Multiphoton fluorescence (MPF) and second harmonic generation (SHG), are particularly useful in highlighting tissue structure and function (*3*, *4*). Calcium, voltage, and molecule-sensitive reporters continue to push the boundaries of multiphoton applications in biomedical research. Vibrational spectroscopic contrast using nonlinear Raman imaging (NRI), namely coherent anti-Stokes Raman scattering (CARS) and stimulated Raman scattering (SRS), has shown particularly wide-ranging applicability. These include rapid identification of tumor margins (*5*), high-dimensional structural labeling (*6*), other biological processes such as drug delivery (*7*), neurotransmitter release (*8*), and electrophysiological dynamics (*9*). Such imaging can highlight the molecular organization and biochemical composition of tissue over time– offering a glimpse into sub- and inter-molecular dynamics without using exogeneous labels. Multiplexing functional, biochemical, and biophysical (e.g. temperature) contrast in cells and tissues can offer a unique tool in evaluating and imaging biological and physiological processes that may not be possible with the same degree of exploratory power in separate instruments (*1*).

Nonlinear optical microscopy is inherently suitable for multimodal imaging applications. Integrating the different modalities that fall under nonlinear microscopy can help alleviate unnecessary system complexity while providing complementary information about tissue structure and function. Notably, CARS, SRS, MPF, and SHG are all optical processes that can occur and be detected simultaneously or sequentially by selecting the appropriate filters and detectors for the techniques used. Several groups have demonstrated selective combinations of nonlinear imaging strategies to address a range of biological questions, *in vitro* and *in vivo* (*10*), particularly sensing fast biological processes such as neuronal action potentials and calcium activity with combined SRS and calcium-sensitive MPF (*9*). Integration of SRS with optical coherence tomography (OCT) has been shown to augment nonspecific scattering-based contrast with vibrational specificity to image lipid distributions in excised human adipose tissue (*11*). More recently, researchers have employed four or more nonlinear imaging modalities, including CARS, MPF, SHG, and third harmonic generation for live tissue and intravital imaging towards wound healing and cancer metastasis (*12*, *13*). While most of these demonstrated approaches have been applied towards observing biological processes over the span of multiple hours at the molecular and biochemical levels, they are limited in their ability to observe snapshots of sub-second functional, biochemical, and thermal processes.

The influence of temperature in physiological and biochemical processes – particularly surrounding sample damage and physiological modulation - has recently become an area of prominence across biomedical research disciplines (*14*–*16*). As optical imaging and perturbation technologies continue to advance biomedical research, it becomes critical to consider thermodynamic effects particularly on live specimens albeit practically difficult. A flexible imaging system that incorporates molecular, biochemical, and biophysical information from cells to tissues, in *vitro* to in vivo, would enable the study of rapid and dynamic biophysical processes from multiple perspectives.

Measuring temperatures in biological samples at high resolutions spatially and temporally has proven difficult and continues to be an active area of research. Groups have multiplexed blackbody thermal imaging and fluorescence microscopy at cellular resolutions in *vitro* (*17*, *18*), however water absorption generally obscures the ability to visualize cell morphology with thermal imaging alone. Moreover, optical components that suffice for use between 0.4 and 14 μm wavelengths to encompass visible optical and thermal infrared imaging are not readily available. Finite element heat transfer modeling and Monte Carlo simulations of photon transport in scattering media is often regarded as the benchmark approach for temperature estimation of dynamic photothermal processes in biological samples with high water concentrations (*19*). Several indirect approaches have been demonstrated including temperature-dependent changes in fluorophore emission, and probe beam deflection microscopy (*16*, *19*, *20*). Several indirect approaches have been demonstrated including temperature-dependent changes in fluorophore emission, and probe beam deflection microscopy (*16*, *20*). While point-based methods (utilizing thermocouples, customized miniature sensors, or intrinsic fluorescence of rare-earth doped glass waveguides) are more direct, they sacrifice spatial information obtained with temperature mapping (*21*). Previous work has utilized thermal imaging with laser speckle imaging to study cerebral blood flow changes during different methods of anesthesia (*22*). However, despite the potential utility of combining thermal imaging with multimodal nonlinear microscopy, such a combination has not been previously reported. A system that can combine multimodal nonlinear microscopy with thermal imaging would offer unprecedented flexibility to study biological processes in real time.

The goal of this project is to integrate multiple nonlinear imaging methods with thermal imaging so that biophysical, biochemical, and molecular information from dynamic biological processes may be imaged with micron level spatial resolution and millisecond level temporal resolution. Towards this effort, we present an imaging platform that integrates single-band CARS, SRS, MPF, SHG, and wide field thermal microscopy (ThM) for characterizing tissues at varying time scales. Applications of our imaging system are demonstrated imaging both *in vitro* with neural cell cultures and tissue specimen as well as in vivo with acute craniotomy rat preparations. Our platform offers a novel approach enabling imaging techniques with otherwise incompatible optical paths (due to physical limitations of hardware) to be applied on a common sample. We demonstrate that this platform provides a robust tool for researchers to probe the physiology of dynamic biological systems, through the unique integration of vibrational spectroscopic, functional fluorescence, and thermal contrast.

## System Design

The framework of the Multimodal Advanced Nonlinear and Thermal Imaging System, or MANTIS, is a three-armed imaging turret (Customized Bergamo II Platform, Thorlabs Imaging Research, Sterling, VA, USA) that respectively images a sample with either nonlinear imaging, thermal imaging, or widefield white light reflectance imaging which is reconfigurable to adapt additional measurement approaches (Fig. 1A). The mechanical arrangement of distinct imaging arms allows for modalities with incompatible optical instrumentation (e.g. ultrafast near infrared imaging with endogenous shortwave infrared measurements) to be performed sequentially without disturbing or repositioning the sample. A FLIR SC8300-series high-speed indium-antimonide CCD camera equipped with a 4X germanium imaging objective (FLIR Systems Inc., Nashua, NH, USA | Fig. 1B) is attached to the thermal imaging arm of the microscope. The thermal camera relies on the endogenous black/grey body emission of a sample between 3-5μm wavelength under the assumption of homogenous emissivity to infer sample temperatures.

**Fig. 1:**
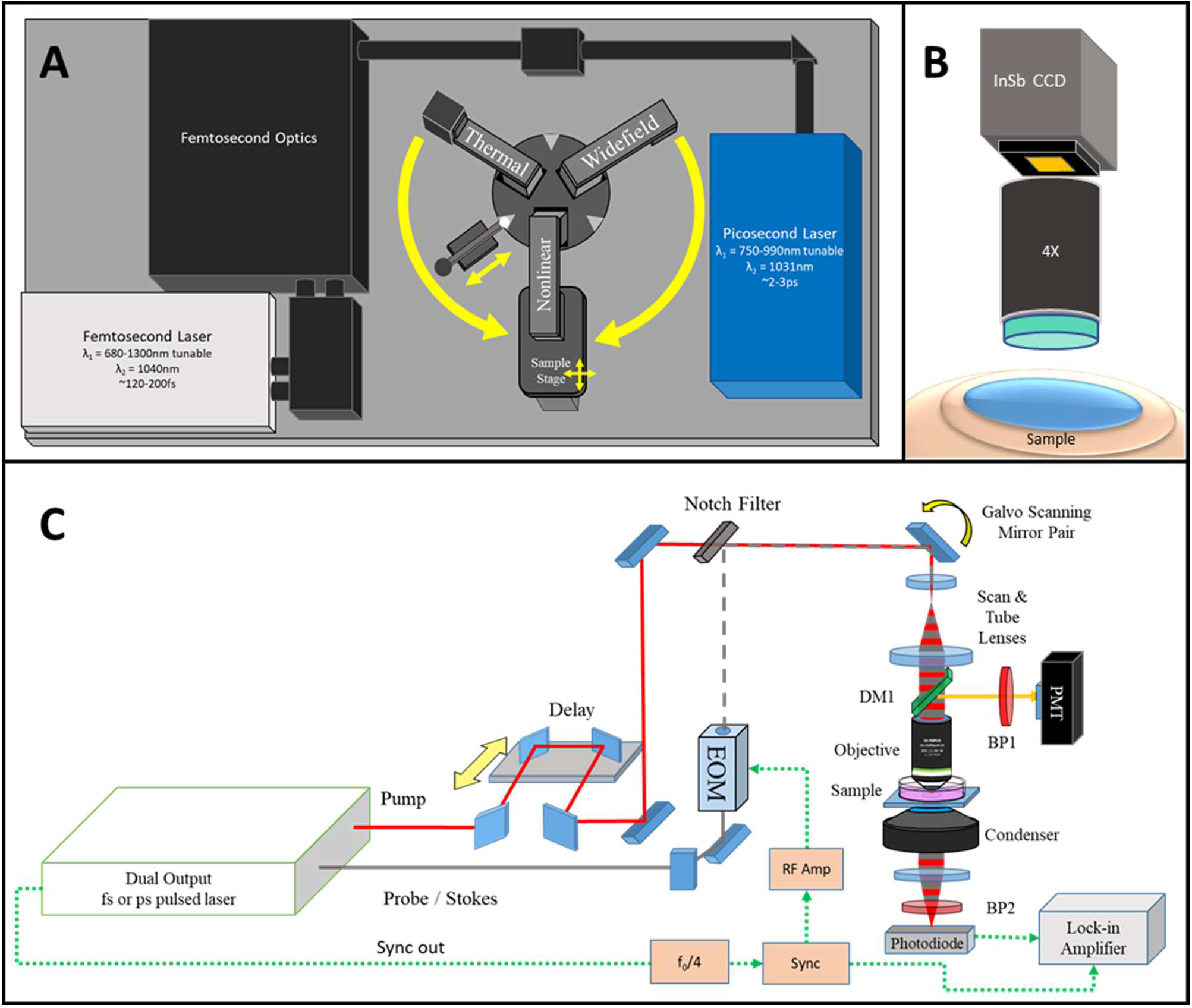
MANTIS System Design. A) The layout of MANTIS. Capable of performing multimodal nonlinear and thermal imaging over the same sample. Additionally, white light imaging on a third imaging arm is available for further expansion of the system. B) Schematic of wide-field thermal microscopy, which infers the temperature of a sample based on the blackbody emission observed between wavelengths of 3 and 5μm. C) General optical schematic of a multimodal nonlinear imaging system capable of integrating CARS, SRS, MPF and SHG imaging. DM – dichroic mirror, BP – bandpass filter, PMT – photomultiplier tube, EOM – electro-optical modulator, InSb CCD – indium antimonide charge coupled device.

The nonlinear imaging section of MANTIS, depicted in Fig. 1C with a simplified schematic, is coupled to two nonlinear laser sources supplying femtosecond (Insight DS+, Spectra Physics, Fremont, CA, USA) and picosecond (picoEmerald S, A.P.E, Berlin, DE) laser lines. Higher spectral bandwidth (~15nm) femtosecond laser pulses provides high peak powers necessary for optimal multiphoton and higher harmonic generation in vivo (*1*). The picosecond laser has narrower bandwidth (~0.5nm, or 10cm^−1^), which is critical for maintaining the spectral resolution necessary for single-band nonlinear Raman imaging while minimizing power at the sample for safe in vivo applications (<20mW average power) (*23*). Both laser systems operate with a pulse repetition rate of 80MHz and output two beams necessary for pump and stokes excitation of NRI contrast processes. The shorter wavelength tunable laser line undergoes a tunable path length delay while the longer wavelength laser line is intensity-modulated at 20MHz via an electro-optical modulator (Thorlabs, Newton, NJ, USA) to facilitate SRS. Each laser’s output is spatially and temporally co-linear and directed with a series of mirrors into the scan head of the nonlinear imaging arm of MANTIS.

The imaging optics of the nonlinear imaging arm are based on conventional upright laser-scanning microscopy. The scanning optics consist of a pair of galvanometric mirrors (Thorlabs, Newton, NJ, USA), which are imaged onto the back focal plane of a commercial objective lens (Olympus XLUMPLFLN, 20X 1.0NA | Nikon CFI Apochromat NIR 60X, 1.0NA) via a 4f optical relay. This scan relay performs a fixed 4X magnification of the laser beam diameter to accommodate the back-aperture pupil size of the largest objective lenses we use relative to the entrance beam diameter (SL50-2P2 & TL200-2P2, Thorlabs, Newton, NJ, USA). Two epi-detection ports with a photomultiplier tube (CARS/MPF/SHG | GaAsP Amplified PMT, Thorlabs, USA) and a large area 50-V reverse-biased photodiode (SRS | A.P.E. GmbH, Berlin, DE) are used for imaging epi-detected contrast. Two additional detachable forward detection ports were also built to accommodate coherent imaging modalities in transparent samples in transmission mode. Stimulated Raman loss for SRS was demodulated via a commercial lock-in-amplifier (A.P.E. Gmbh, Berlin, DE). All emission filters and excitation wavelengths included with the system are summarized in Table S1 (Semrock, Brattleboro, VT, USA). Scanning and detection hardware for imaging is controlled through ThorImageLS version 2.1, 3.0, or 3.2 (Thorlabs Imaging Research, Sterling, VA, USA). All images shown are raw with linear intensity rescaling, with all analysis being performed in FIJI (*24*).

The third imaging arm of MANTIS initially features a color complementary metal oxide semiconductor (CMOS) camera (Thorlabs Inc., Newton, NJ, USA) with a variable focal length lens (Navitar Inc., Rochester, NY, USA) for widefield white-light reflectance imaging of samples (Fig. 1A). However, this arm was built into the imaging system to readily enable integration of additional contrast modalities that may be further incompatible with multimodal nonlinear microscopy – such as interferometric contrast or stereoscopic surgical guidance. Each arm is locked into position by a custom-designed spring-loaded locking system against the rotating turret base (Fig. 1A). The maximum fields of view of each imaging arm is ~800μm-x-800μm for nonlinear microscopy with a 20X objective, 3mm-x-4mm for thermal microscopy with a 4X objective, and 50mm-x-50mm for widefield white light reflectance imaging.

## System Performance

To achieve high-speed imaging at subcellular resolution, MANTIS was designed to resolve at least 1-μm lateral resolution in the nonlinear imaging arm. Performance was verified on a series of standards, (Fig. **2**), with measured axial and lateral resolutions for each modality summarized in Table S2 (data summarized in Table S1). Images of 0.5-μm diameter polystyrene (PS, latex, Polysciences Inc. Warrington, PA, USA) beads with CARS and SRS were obtained to generate a point spread function (PSF), which is reported here as the full-width half-maximum intensity cross-section of beads fit to a Gaussian curve. Fluorescent bead calibration standards with a 250-nm diameter (Polysciences Inc. Warrington, PA, USA) were used to measure the PSF of MPF. All PSF calculations were performed using the MetroloJ plugin in FIJI (*25*). Measured resolutions, reported in Table S2, are on par with results published literature when imaging with numerical apertures approaching 1.0 (*3*, *23*, *26*). Spectral separation of polymethyl-methacrylate (PMMA, acrylic, Polysciences Inc. Warrington, PA, USA)) and PS beads with single-band SRS are shown in Fig. **2**B. Acrylic (PMMA) beads are depicted using the 2927-cm^−1^ band – an asymmetric CH3 resonance- (yellow), and latex (PS) beads are depicted spectrally separate with the 3053-cm^−1^ band− an asymmetric CH_2_ resonance- in blue. The signal-to-noise ratio (SNR), measured at the edge of a vegetable oil meniscus at the 2927-cm^−1^ band with SRS (Fig. S2), was calculated to be 34.6 (*27*).

**Fig. 2:**
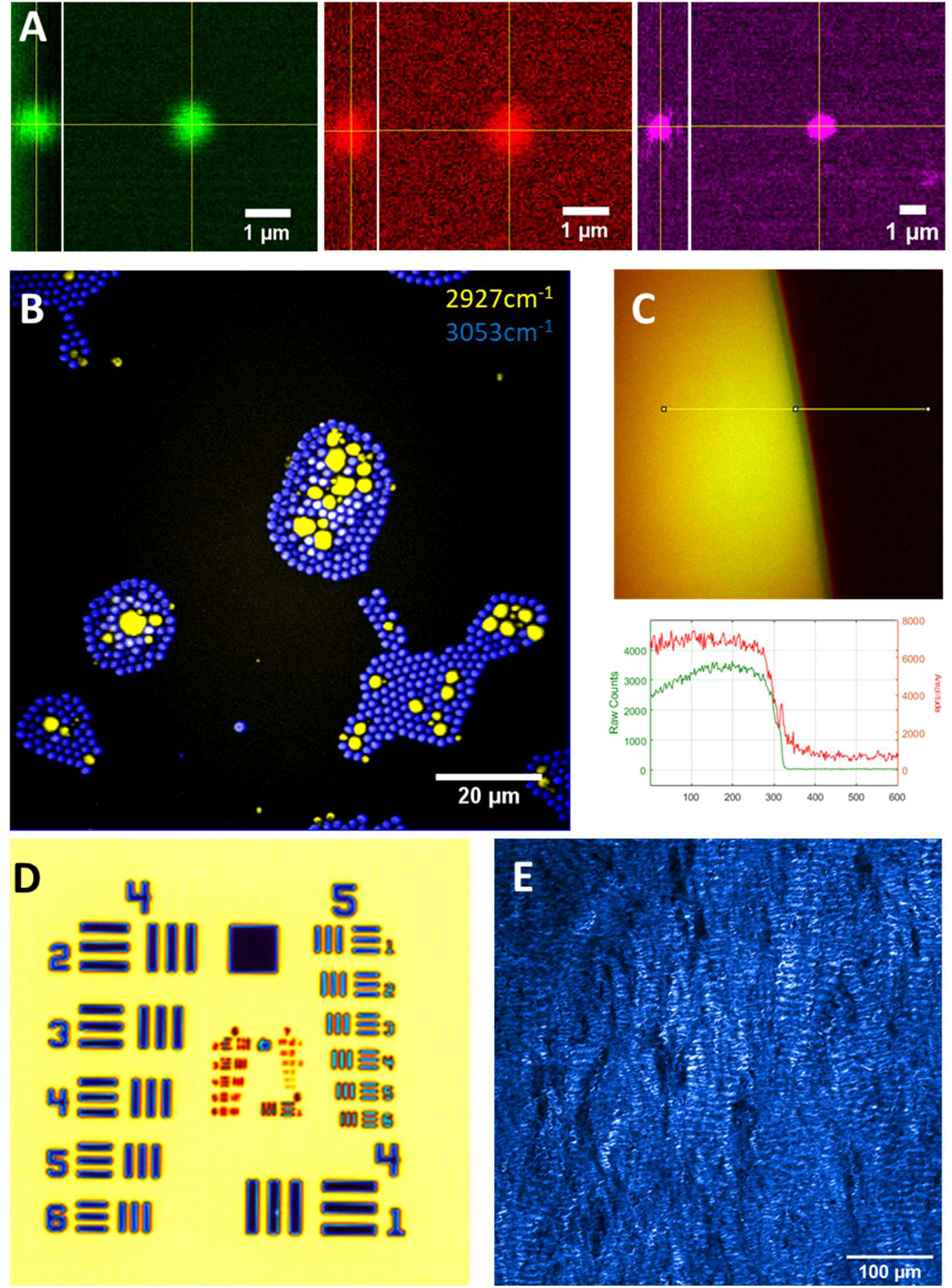
MANTIS System Performance. A) Resolution of CARS (Green), SRS (Red) with a 1μm polystyrene bead, and Multiphoton Fluorescence (Magenta) with a 500-nm yellow-fluorescent polystyrene bead. Measured resolutions summarized in Table S1. B) Preparation of mixed beads containing 2μm polystyrene beads (blue, 2927cm^−1^) with 1-10μm PMMA beads (yellow, 3053cm^−1^). Peaks specific to each bead type C) CARS (Green) and SRS (Red) of signal-to-noise profile measured at a vegetable oil – air meniscus at 2927cm^−1^. D) Thermal Imaging of a 1951 U.S. Air Force Target. E) SHG imaging of a porcine mitral valve ex vivo.

Resolution standards to verify SHG imaging resolution are not commercially available. In place of a controlled standard, a collagen-rich biological sample (a porcine mitral valve) was imaged at a high spatial sampling density (250-nm/px) and the finest resolvable fibrillar structures were measured as a proxy for lateral resolution. Sample images of mitral valve collagen are shown in Fig. **2**E and Fig. S3. Example calculations of fibril diameter are show in Fig. S1. Fibrillar collagen bundles and quaternary structure are clearly visualized. While fibrillar structures vary substantially in diameter (*3*), the resolving power of SHG on MANTIS (Table 2) is well within our targeted resolution goals being able to resolve sub-micron spatial features.

The FLIR thermal camera resolution was estimated by identifying the smallest resolvable group on a 1951 United States Air Force resolution target. Resolving group 6, element 4 equates to a lateral resolution of 6.9-μm. The depth of focus of the thermal camera is about 40-μm, based on an f/4.0 aperture stop of the objective lens in the thermal camera. Since the depth of focus of the thermal camera is substantially larger than the depth of focus of the nonlinear imaging system, registration in the axial dimension straightforward. The factory measured temperature resolution is specified to be 0.1°C. The nonlinear and thermal microscopy field view were independently adjusted to centrally registered the corner of a 10-um grid target. Switching between thermal and nonlinear microscopy fields of view was found to repeatable within 1-μm radial to imaging turret rotation and 10-μm tangent to imaging turret rotation (Fig. 1A, yellow arrows). Since field of view of the thermal camera is nearly four times that of the nonlinear microscopy field of view – making the coincidence of each modality’s field of view with each other straight-forward Figure S5. Once aligned, the fields of view were observed to remain well registered for more than a week following initial alignment. It takes about 90-180 seconds to switch between nonlinear and thermal imaging arms over the same sample. The sub-millimeter repeatability of positioning multiple imaging arms readily enables multiplexing of thermal and nonlinear microscopy on the same field of view without needing to reposition the sample.

## *In vitro* Imaging

To verify the capability of MANTIS to perform in *vitro* imaging, cultured NG108 cells – a spiking neuron-astrocytoma hybridoma cell line - were imaged with CARS and MPF, as seen in Fig. **3**. Briefly, cells were plated on poly-D-lysine coated cover glass for 48hr prior to imaging and maintained in Dulbecco’s Modified Eagle Medium (DMEM) supplemented with 10%v/v fetal bovine serum and 5mM L-glutamine. 12-hours prior to imaging, medium was replaced with DMEM containing 3%v/v FBS to morphologically differentiate adherent cells. Performing CARS imaging at 2830-cm^−1^ resonance highlights a lipid-dominant CH_2_ symmetric stretch mode, which relates to cellular projections and lipid-rich intracellular contents such as organelles, lipid droplets, and vesicles (Fig. **3**A). Autofluorescence from NADH and FAD can be used to extrapolate metabolic information relating to relative aerobic metabolism dynamics and metabolic cofactor distribution throughout the cell, Together, combined CARS and MPF demonstrate the structural and functional capacities of multimodal nonlinear imaging in *vitro* (Fig. **3**B and C). Blackbody thermal imaging of cells in media in an upright configuration was not feasible due to the strong absorption of water in the wavelength range measured by the thermal camera (3 to 5-μm). Imaging with cellular resolution with thermal microscopy is possible in an upright configuration (Figure S4A), with the ability to resolve cellular temperature distribution. However, once medium is introduced to the imaging field of view (Figure S4B), cellular morphology is occluded due to strong water absorption of the sensed wavelength range. This has been addressed by others by utilizing inverted imaging configurations (*17*, *28*, *29*), however resolvable cellular morphology were not demonstrated.

**Fig. 3:**
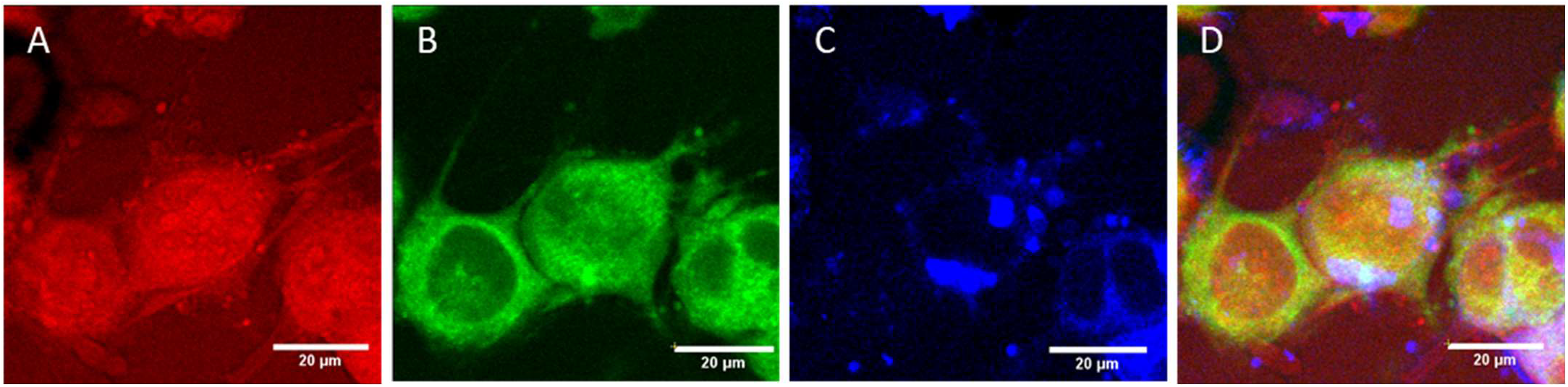
*in vitro* imaging capabilities of MANTIS. NG108 cells observed with CARS (A) at 2830cm^−1^, a protein-dominant peak, NADH autofluorescence (B), FAD autofluorescence (C), and as a composite overlay of the three channels (D). Images were acquired separately and coregistered together with a rigid transformation.

Combined SHG and MPF was demonstrated on fresh *ex vivo* porcine mitral valve samples (Figure S5). Tissue samples were harvested from porcine cardiac tissue postmortem and stored in phosphate buffered saline for 72hrs prior to imaging. Samples were mounted on a standard microscopy slide and imaged through leveled and supported cover glass in contact with the tissue surface. Simultaneous autofluorescence and SHG under 930-nm illumination with the femtosecond laser source yielded high resolution microstructure of endogenous elastin and collagen, free of exogenous labeling (*30*). Changes in collagen and elastin can dangerously disrupt the structure and function of heart valves (*31*). Having methods to study new interventions to improve heart valve function could be valuable to cardiac biomechanics researchers. Combining SHG with SRS at 2880-cm^−1^ to broadly highlight lipids allows for the visualization of tissue architecture in unfixed frozen sections of murine cervix (Figure S6). Distinct chemical and structural differences are notable between the epithelium and stroma of the cervix. The stroma is dense with collagen and low in cellular content, while the epithelium is rich in lipids and proteins due to cellularization. The lamina propria of blood vessels are also discernable with SHG, appearing as fine tubular structures with strong SHG signal, providing an indication of tissue vascularization. Visualizing the changes in vasculature, cellular, extracellular matrix protein distribution throughout cervix shows potential for studying parturition processes and cancer progression in the cervix without the need for exogenous labelling.

Fig. **4**, Figure S6, and Figure S7 demonstrate multimodal nonlinear imaging with CARS, SRS, SHG, and MPF in an ex vivo rat sciatic nerve preparation. Nerves were harvested from Sprague-Dawley rats and imaged immediately postmortem. Contrast from CARS and SRS primarily highlights myelin (Figure S7B&C). The CARS and SRS images correspond well with the MPF images of FluoroMyelin green (ThermoFisher Inc | Figure S7E, Figure S8C and D) but are less prone to heterogeneous dye uptake. While myelin plays a major functional role in nerve signal conduction, collagen is a major structural component of the sciatic nerve that provides a protective exterior sheath and additional mechanical stability within the nerve. Collagen contrast from SHG can be used to visualize the epineurium and fascicle-residing collagen (Figure S9). These four modalities together therefore offer key structural insight to the sciatic nerve and can be used to study nerve injury, regeneration, and disease. Furthermore, the ability to flexibly apply these techniques to different sample types, between live cells and excised tissue specimen, highlights a key design goal of the imaging platform.

**Fig. 4:**
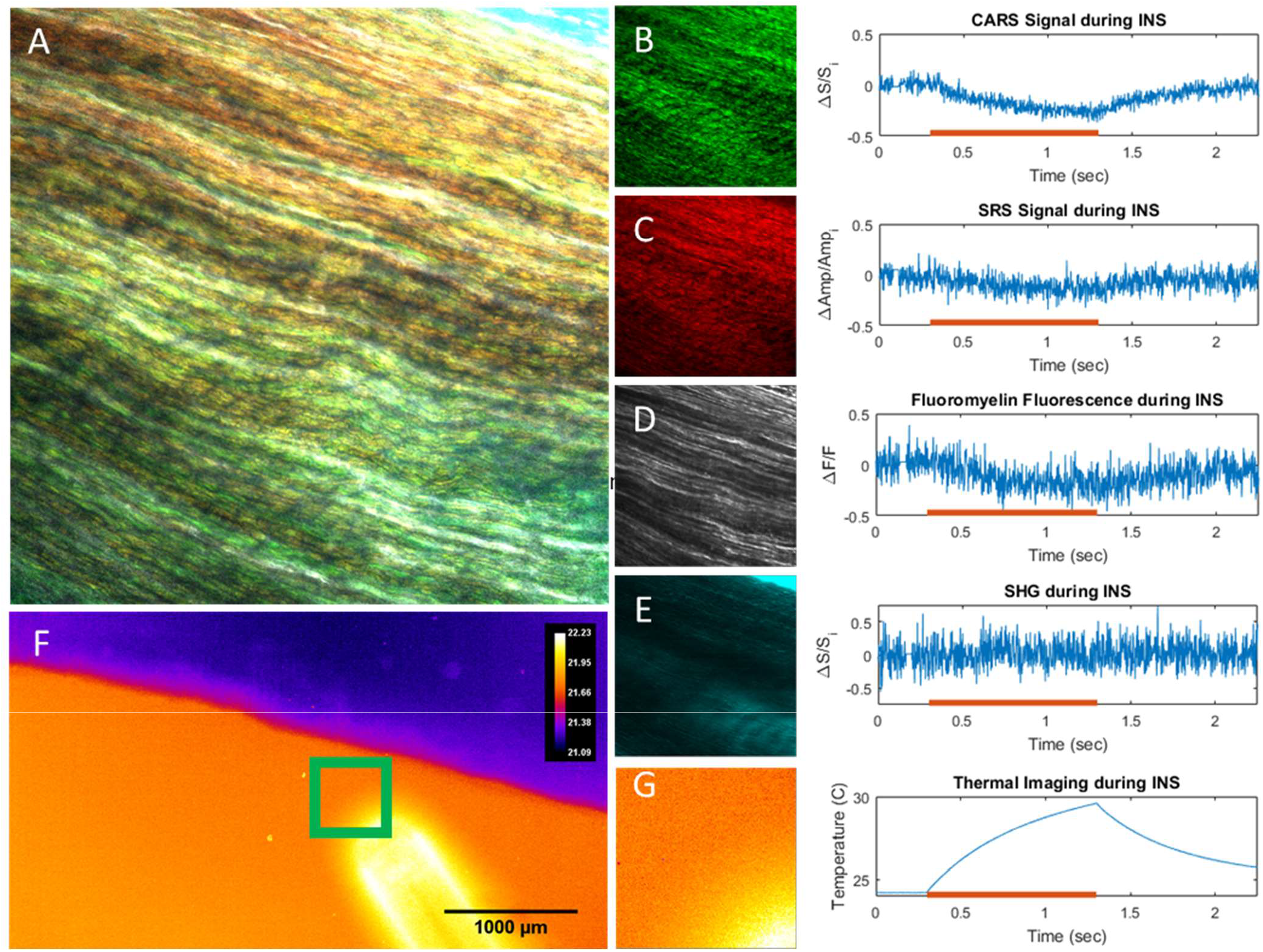
Ex vivo rat sciatic nerve samples imaged with high-speed multimodal nonlinear and thermal microscopy. A) Combined image of B-E. Images and corresponding high-speed line scans through the centers of the fields of view with B) CARS at 2880-cm^−1^. C) SRS at 2880cm^−1^, D) FluoroMyelin Green, E) SHG. Red lines on graphs indicate the exposure of the tissue to 1875nm light from an optical fiber. Light exposure causes a local temperature increase over 1 second (thermal images, F&G) yielding a corresponding decrease in nonlinear signals.

The key advancement allowed by the MANTIS platform is the ability to multiplex optically incompatible imaging techniques such as thermal imaging with nonlinear optical microscopy. Thermal imaging can be used to evaluate fast photothermal processes, such as infrared neural stimulation (*32*, *33*), with coregistered contrast from nonlinear imaging modalities to study real time biophysical dynamics. To assess the multimodal capabilities of MANTIS, readout of multimodal nonlinear contrast was directly correlated with high-speed temperature changes during infrared neural stimulation in a fresh ex vivo sciatic nerve preparation (Fig. **4**). Point-scanned line repeats were employed to obtain high framerate nonlinear signals during IR stimulation. Intriguingly, our results show a notable decrease in nonlinear signals that tracks closely with temperature increases over the period of one second. Notably, SHG signals did not exhibit a discernable decrease in signal, which is attributable to the lack of collagen present through where line scans were acquired (center of the field of view, Fig. **4**). However, decreases in SHG signal during IR stimulation have been verified in other FOVs, suggesting this signal decrease is a more general physical phenomenon. Decreases in nonlinear signal were found to be caused by a defocusing artifact induced by the spatial thermal gradient inherent to infrared neural stimulation in the microscope’s field of view. Resolving photothermal effects of IR heating in biological tissues at high speed, such as during INS, will provide invaluable information in studying in vivo applications of INS.

## *In vivo* Imaging

An acute craniotomy preparation in an anesthetized rat model was utilized to validate the in vivo capabilities of MANTIS. All animal use protocols were approved by Vanderbilt University Institute for Animal Care and Use Committee (VU-IACUC). Sprague-Dawley rats were anesthetized with intraperitoneal injections of ketamine (40-80mg/kg) and xylazine (10mg/kg) for 40 minutes prior to surgery. Anesthetic depth was monitored every 10 minutes and maintained with follow-up half-dose anesthetic injections every 90-120 minutes as needed. Animals were mounted in a stereotaxic frame under the microscope objective while a 2mm-x-2mm section of skull and dura mater overlying the somatosensory cortex was carefully removed. Tissue hydration was maintained with sterile saline throughout imaging. Animals were sacrificed by anesthetic overdose and cervical dislocation following experiments. During imaging experiments, pixel dwell times needed to be increased to 8-μs to account for signal loss in moving the SRS detection path into an epi-detection configuration. Epi-SRS detection was implemented by replacing the dichroic mirror immediately preceding to the objective in the incident light path with a polarizing beam splitter cube and quarter wave plate (*23*). Images were initially acquired at a high spatial sampling density (0.5-μm/px, 800×800 px) with a 0.2Hz framerate. However, reducing the spatial sampling density and overall image sizes (3μm/px, 256×256 px) over a comparable field of view, we were able to achieve 2Hz framerates. Subsequently, repeated point-based line scanning could be implemented with a 488Hz line rate to observe faster biophysical dynamics. This flexibility emphasizes the trade-offs between spatial and temporal sampling in nonlinear microscopy approaches in vivo. Acquiring images using CARS, MPF, or SHG needed to be performed sequentially, due to the availability of only one PMT detector at the time of experiments. However, any one of the three PMT-requiring modalities can be multiplexed with SRS contrast, since SRS uses a separate detector in an adjacent detection path. Excitation and detection wavelengths are summarized in Table S1.

Examples of in vivo images can be seen in Fig. **5**. A lipid-rich CARS resonance at 2850cm^−1^ (Fig. **5**C) and a protein-dominant SRS resonance at 2930cm^−1^ (Fig. **5**B) appears to highlight superficial cellular features as well as vascular perfusion within blood vessels. Autofluorescence contrast from blue and green detection channels are most commonly used to measure endogenous NADH (Fig. **5**G, Table S1) and FAD (Fig. **5**E, Table S1). This diffuse NADH signal is likely arising from the neuropil at the pial surface of the cortex, while FAD tends to present more sparsely superficially in the outer cortex in vivo; this weak fluorescence is often spectrally indistinct from other biological autofluorescence (*34*, *35*). Contrast from SHG (**Fig. 5** revealed unknown morphologies, which may be due to cytoskeletal structures across the pial surface, but is more likely to be residual collagen from the dura mater after surgical preparation. Thermal imaging (**Fig. 5H**) of the brain surface appears to reveal mostly topographical contrast - vasculature is clearly visible. The lack of optical penetration depth of measured wavelengths of the thermal camera are likely responsible for the brain surface topographical contrast. Temperature fluctuations across the surface of the brain appear to be minimal, even with breathing, heart rate, and motion artifacts. Larger vessels appear slightly warmer than smaller vessels. Focusing artifacts appeared to have some effect on the accurate approximation of the brain surface temperature within 1°C – particularly noticeable at the edges of the field of view, which likely arise from out of focus signal from underlying bulk tissue due to the high degree of surface curvature in the sample (**Fig. 5H**). Cellular morphologies were not apparent in thermal images of the brain surface. Thermal imaging of the brain surface was capable of acquisition speeds approaching 180Hz framerates with a full field of view. This imaging rate can be substantially increased to 500Hz or more by cropping the image sensor readout without loss in spatial resolution or pixel binning without reduction in field of view. The vessel morphologies visible between thermal and CARS/SRS contrast made fine registration of thermal and nonlinear imaging fields straightforward with rigid registration of manually labelled image features.

**Fig. 5:**
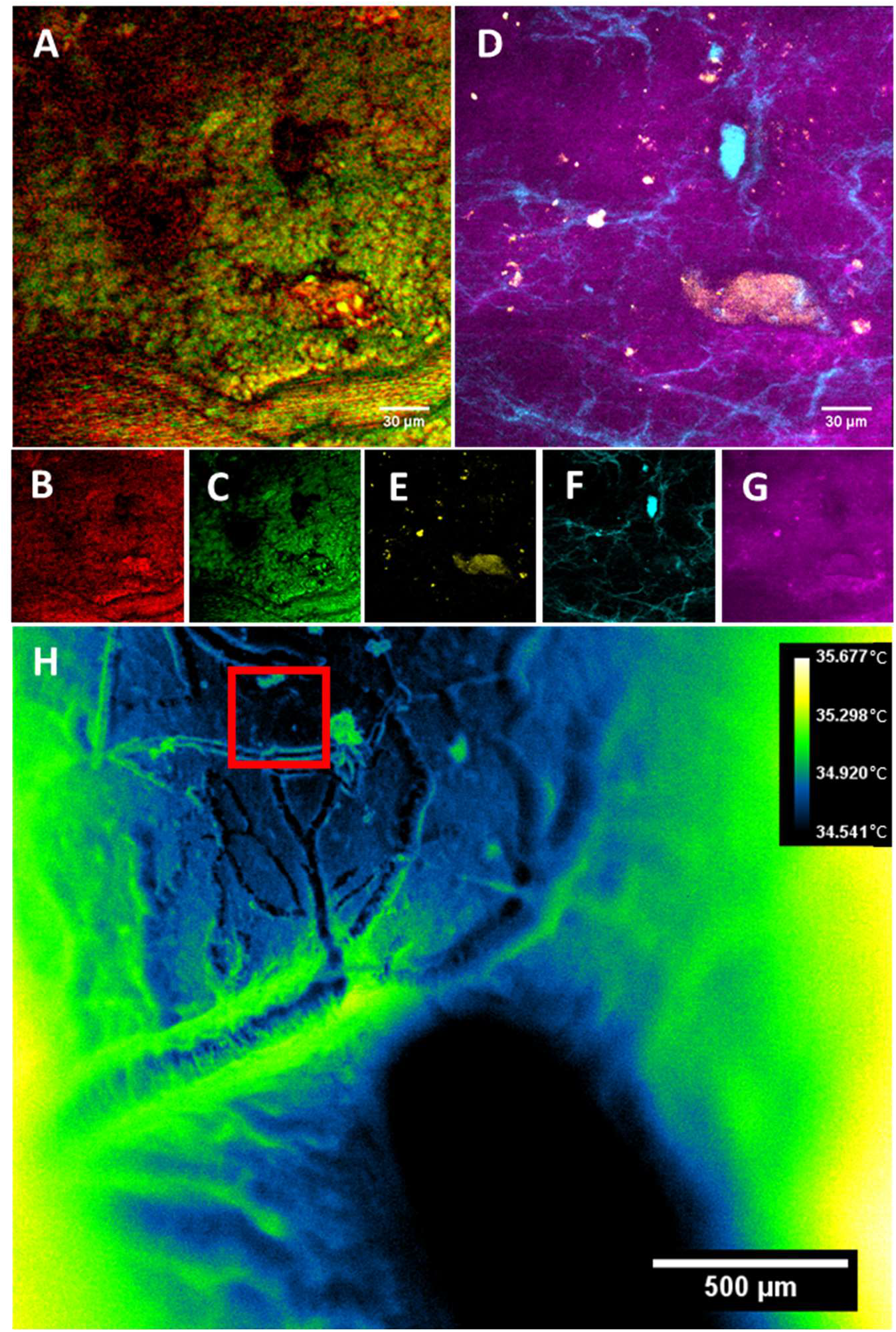
Multimodal images from a rat brain in vivo. A) Composite image of B (SRS, 2930cm^−1^) and C (CARS, 2850cm^−1^) highlighting lipid and protein-dominated signals, respectively. D) Composite image of the same field of view in A of E (Green autofluorescence), F (SHG signal), and G (Blue autofluorescence, likely NADH). (H) Thermal images of the brain surface, temperatures shown in °C.

## Combined Multimodal Nonlinear and Thermal Microscopy

Infrared neural stimulation (INS) utilizes pulsed short-wave infrared laser light to transiently invoke a thermal gradient in neural tissue resulting in activation of neural cells. One of the major concerns about neuromodulation with rapid targeted thermal gradients is that the change in temperature could cause cell damage. Cells are naturally prone to damage at elevated temperatures; however, the role of temperature-time history is often underappreciated in the context of biological thermodynamics. Integrating fast nonlinear and thermal microscopy offers a particularly useful platform for studying the physical and functional impacts of INS on neural cells.

The ability to correlate high resolution temporal and spatial thermal information (Fig. 6E&F) with the functional information afforded by nonlinear imaging (Fig. 6A-D), available in the MANTIS platform offers a unique opportunity to visualize biochemical, biophysical, and biomolecular dynamics of cultured NG108 cells during INS. An 8ms pulse of 1875nm infrared light was delivered to the cells via a 400μm diameter optical fiber at average radiant exposures spanning 0.5 (stimulating) and 3J/cm^2^ (damaging) while imaging at 10Hz framerates. Multiphoton fluorescence of cells imaged in saline containing a damage indicator, propidium iodide (PI, 1uM concentration, Fig. 6C, Table S2), differentiated healthy and damaged cells due to IR exposure. Increases in relative fluorescence greater than 3% were presumed to be indicative of cell damage. Simultaneously, SRS imaging at a CH3 resonance (2930cm^−1^, lipid/protein) provides endogenous biochemical information of cells during INS. In this study, SRS images were primarily used to segment cell morphologies to extract viability status (MPF) and interpolate thermodynamics (thermal imaging).

**Fig. 6:**
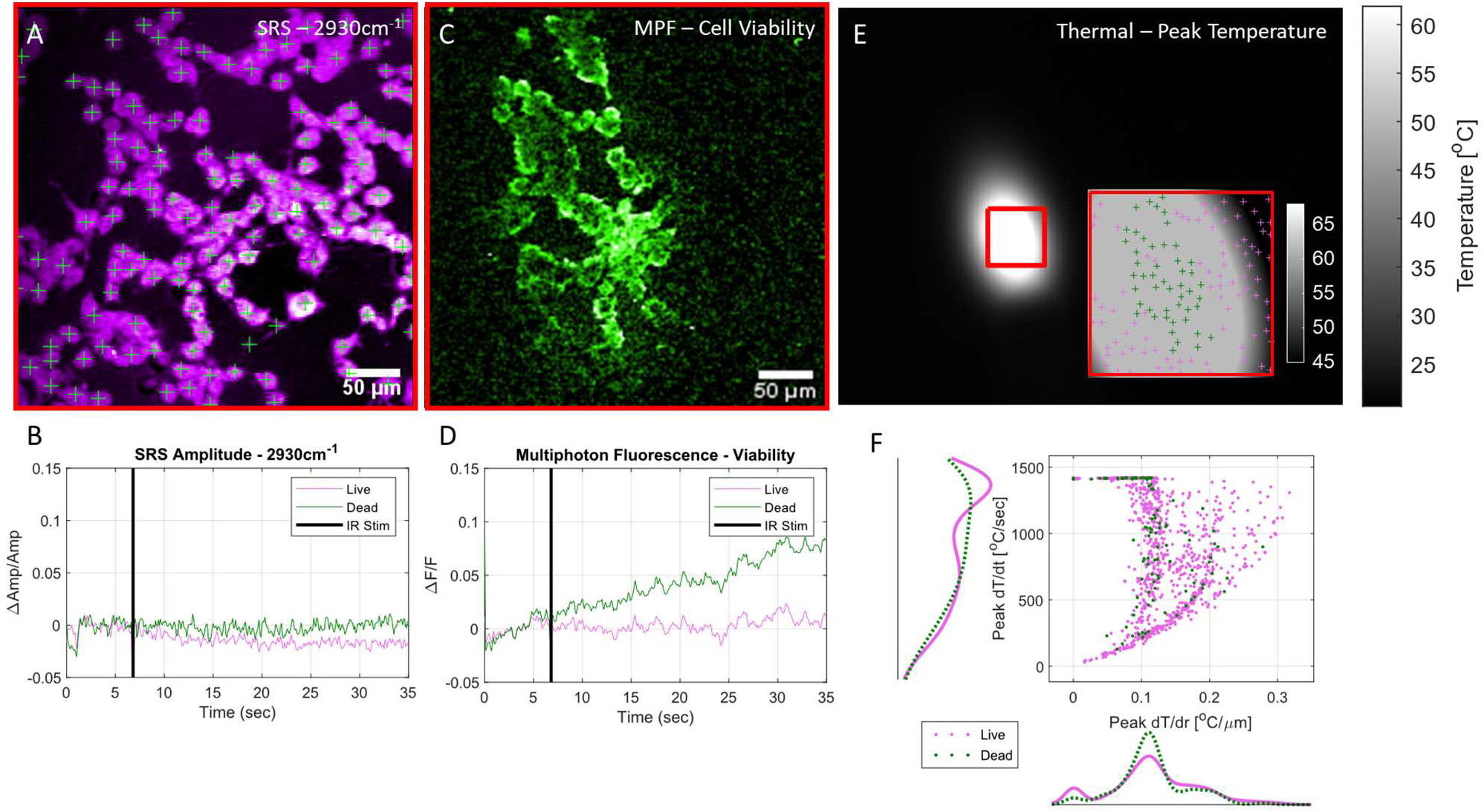
Fast Multimodal Nonlinear and Thermal Microscopy of NG108 cells during in infrared neural stimulation –. Examples of an A) SRS average intensity projection image of cells with indicated centroids and (B) pooled average SRS timeseries of live (magenta) and damaged (green) cells during IR stimulation across all observed cells. (C) Example of a normalized sum intensity projection of cell viability image timeseries used to indicate cell viability via uptake and fluorescence of propidium iodide. (D) Average pooled MPF timeseries of all observed live and damaged cells during INS. Increases in MPF signal after INS above 3% dF/F over 30 seconds were considered damaged/dead. (E) Thermal images of NG108 sample preparation during peak temperatures of INS. Registration of thermal and nonlinear fields of view (E, inset) allows for precise spatial temperature mapping. Cellular details are not observable due to strong absorption by the aqueous imaging medium. Coupled with high speed thermal imaging, (F) high resolution temporal and spatial thermal observations can be directly correlated with functional cellular outcomes such as cell viability. Data shows results of 10 different experiments including 3 different INS intensity conditions, n = 1144 cells.

Thermal images during INS were acquired at 34Hz framerates after all nonlinear imaging and stimulation experiments were completed. Temperature information is not simultaneously available with nonlinear observations due to physical limitations. Since the thermal properties of cells mounted in imaging saline behave thermodynamically like water upon INS, post-hoc registration of thermal information with *a priori* knowledge of thermal and nonlinear fields of view was used. Multimodal image registration was performed with an iterative closest point algorithm implemented in MATLAB to register manually selected features of a fluorescent target (Chroma, Rochester, NY, USA) observable in both nonlinear and thermal imaging modes. Cell centroids were calculated from cell morphologies identified with SRS (Fig. 6A) in FIJI utilizing a seeded watershed segmentation algorithm (*24*, *36*). Spatial transformations from aforementioned field of view registration were applied to the cell centroid positions to obtain their positions in the thermal camera field of view. Cell positions in the thermal camera field of view were used to interpolate temperature-time and temperature-space thermal data on a per-cell basis utilizing built-in 2D interpolation functions in MATLAB.

Since the stimulating IR light (1875nm) is strongly absorbed by water, the immersion medium required for nonlinear imaging had a substantial impact on the heating of cells, leading to potentially large discrepancies in thermal imaging measurements. To minimize optical absorption of 1875nm stimulation light by the aqueous immersion medium necessary for the nonlinear microscopy objective (Olympus XLUMPLNFL 20X, 1.0NA), spectroscopic-grade deuterated water (Sigma Aldrich, St. Louis, MO, USA) was used as an immersion medium (*33*). Cells remained immersed in normal imaging media and were separated from heavy water by the coverslip on which the cells were mounted. The reduction in IR absorption in the objective lens immersion media was enough to allow for representative spatial thermal measurements of INS on aqueous cellular samples with the thermal camera. Functional observations of cell death aligned well with peak spatial temperature maps (Fig. 6E, inset). It is expected that absolute temperature measurements at the sample are likely to differ with and without the presence of deuterated immersion medium during INS, however the spatial and temporal dynamics of IR-induced cellular thermal gradients are expected to remain consistent and comparable across all exposure conditions. With this assumption, physiological comparisons can be effectively drawn with the understanding that the observed absolute temperatures are likely slightly higher than during nonlinear image experiments where immersion medium may impact INS.

Spatially registered observations of cell functional, chemical, and temperature dynamics is only made possible by a combined and accurately coregistered nonlinear and thermal imaging platform such as the MANTIS platform. From these multimodal imaging experiments, we observed that cells that experience more rapid changes in temperatures as a function of time were more likely to be damaged (Fig. 6E, F). Damaged cells are more likely to be located near low spatial thermal gradient values, which correspond to local maxima in spatial heating profiles (Fig. 6E, inset). However, cells outside the fiber illumination would be expected to present similar spatial thermal gradients as cells at peak levels of heating - introducing some heterogeneity in the spatial thermal gradient information. Considering spatial and temporal thermal gradient information together clarifies the thermodynamic difference between heated and unheated cells. Numerous cells appear to survive rapid temperature changes while others do not, however damaged cells are far more likely to undergo rapid heating. This observation demonstrates the variability in cell physiological responses due to INS. Distinct increases in mean multiphoton fluorescence timeseries indicate cell damage within 30 seconds of IR exposure. Cell damage appears to coincide with increases in mean SRS CH3 signal following IR stimulation. The basis for SRS signal increase is currently unclear, though we speculate attribution to endogenous cellular damage responses (e.g. lipid vacuolization, organelle damage, and increased chaperone protein expression such as heat shock proteins). There remains to be extensive validation of changes in endogenous SRS signal in the context of live cell imaging and damage beyond lipid storage and cell membrane dynamics. Experimentally, SRS images were primarily used in Fig. 6 to identify, segment, and locate cells for temperature and viability data calculations. However, this platform readily enables such explorations into the molecular basis of SRS signal changes simultaneously with more established live cell fluorescence imaging probes.

## Discussion

We demonstrate the first imaging system that combines CARS, SRS, MPF, SHG, and thermal imaging into a single microscope for biomedical imaging applications. The similarities in illumination and detection instrumentation needed for nonlinear excitation has been exploited in the past to design multimodal imaging platforms. But as demonstrated here, our novel system design includes a movable turret that allows for overlaying these different modalities to achieve a more complete picture of biological processes to be imaged. This logically suggests that another imaging modality that typically could not be integrated into a nonlinear optical microscopy path may be similarly incorporated. Such flexibility of optical microscopy design provides the opportunity to study physiology in unique and dynamic ways.

The proposed imaging system, MANTIS, was built with modularity and expandability in mind. On the nonlinear imaging arm, the current configuration has four optional detection ports - two each in the epi- and forward detection configurations. Additional channels can easily be added to expand simultaneous imaging capabilities based on research needs. Lock-in detection arms can be refitted with the appropriate optical filters for transient absorption imaging. Sum-frequency generation imaging can also be readily integrated for studying ordered molecular orientation and interfacial phenomena in biological samples. The widefield reflectance imaging arm may be useful for integrating laser speckle, diffuse reflectance, optical coherence tomography and microscopy, or spatial frequency domain imaging with the correct illumination optics to map tissue blood flow, oxygenation, and optical properties. Furthermore, the widefield reflectance and thermal imaging arm were built on detachable 96-mm optical rails so that other imaging methods may be integrated based on future research needs. The multi-armed imaging concept can also yield more utility out of a condensed instrument footprint, which may be useful in places where laboratory floorspace is limited.

The scanning optics used in nonlinear imaging arm can achieve 1kHz scan rates. A galvo-scanning pair was chosen over a faster galvo-resonant scanning pair, which typically offer an order of magnitude increase in scanning rates. By doing so, improved control over image sampling densities can be achieved to readily span subcellular and multicellular scales. This is exemplified by the difference in sampling density observed between Fig. **3** and Fig. **5**. As demonstrated in Fig. **4**, our imaging system can perform nonlinear imaging with line scans approaching 0.5kHz with detectable amounts of signal. This is enough to resolve high-speed biophysical dynamics, such as during neural modulation or optogenetic stimulation, from a functional, chemical, and physical standpoint. Kilohertz bandwidth scanning rates are sufficient to achieve visualization of neuronal action potentials as demonstrated by Lee et *al*. with balanced detection SRS and calcium fluorescence microscopy (*9*).

Both femtosecond and picosecond lasers were included in the design of MANTIS for signal and spectral resolution considerations. The bandwidth of the picosecond laser source (around 0.5nm) allows for higher spectral resolutions (around 10cm^−1^) which is critical for nonlinear Raman imaging (*37*). However, the peak powers of the picosecond source are orders of magnitude lower than that of the femtosecond laser, yielding less signal during multiphoton fluorescence and higher harmonic imaging. At the time of construction, the use of broadband nonlinear Raman techniques, such as pulse shaping and spectral focusing had yet to be realized for fast imaging in vivo (*38*). As such, design considerations to optimize narrow band spectral resolution and in vivo imaging speed were a priority. A number of broadband spectral NRI techniques have since been described (*39*, *40*), with a handful demonstrated in vivo (*10*). The flexibility of the existing instrumentation on MANTIS readily allows for the integration such broadband approaches.

Using blackbody thermal microscopy to characterize sample temperatures in conjunction with nonlinear imaging, or any laser-scanning microscopy approach, has yet to be previously published. Estimating temperature with a thermal camera provides a more direct measurement than other approaches, such as fluorescence-based methods. The ability to measure thermal information alongside the functional and structural information offered by nonlinear microscopy provides a unique instrument to explore new questions in biophysics, particularly ex vivo and in vivo. Practically, thermally microscopy presents some advantages and drawbacks depending on the model systems being imaged. We have found that high resolution fluorescence microscopy can be difficult to interpret in highly thermally dynamic systems with high numerical apertures at high imaging speeds, which became apparent in applying infrared neural stimulation in sciatic nerve (Fig. **4**) as well as in cells. Having thermal information to contrast and systematically compensate for thermally induced effects is something that our imaging platform lends itself to accomplish. Since imaging depth with thermal microscopy is limited due to the optical penetration depth of water in the 3-5μm wavelength spectral regime, this means that SWIR based imaging methods are ideally suited for measuring surface temperatures in water-dominant samples such as biological tissues. This is even more apparent in vitro, where imaging through any amount of aqueous medium occludes cellular morphology (Figure S4). While others have demonstrated blackbody thermal imaging of cells via inverted microscopy through cover glass (*17*, *18*), visualizing cell morphology with thermal imaging has not been previously demonstrated. Thermal microscopy on MANTIS can be performed readily and reliably on tissues ex vivo (Fig. **4**F) and in vivo (Fig. **5**H). Topographical features such as blood vessels tissue surfaces are clearly visible. The pronounced topography of these anatomical features is helpful when performing fine registration nonlinear and thermal imaging fields. Temperature at aqueous interfaces are useful in approximating spatial temperature distributions in vitro. Combining surface temperatures with finite element heat transfer modeling and precise control over sample thermodynamics can be used to render volumetric estimates of temperature in vitro (*18*). In combination with fluorescence or other nonlinear imaging modalities, imaging the thermodynamics of adherent cells in vitro or in tissues with thermal microscopy could be quite useful. Consequently, multimodal methods almost become an essential requirement to correlate temperature and real-time cellular observations. Our microscopy platform readily allows such observation to be made and flexibly conFig.d to address a wide range of biological and physiological questions.

As expected, cell viability at elevated temporal thermal gradients values is more likely to result in cell death than at lower gradient values (Fig. 6F, axes histograms). However, as evident by the overlap in cell viability as a function of thermodynamics in Fig. 6F, peak thermal gradients are not absolutely predictive (Fig. 6F, scatter plot). While cells heated quickly are more likely to become damaged, rapid heating is not necessarily a death sentence for cells. The variability in thermally evoked cell damage illustrates the need for tools to study underlying functional and biomolecular dynamics evoked by IR light on a cell-to-cell basis. Exposure time and time after exposure becomes a crucial dimension of cellular physiology to explore in the context of infrared neural stimulation. Endogenous lipid/protein signals (Fig. 6A&B) and a fluorescence marker for cell damage (Fig. 6C&D) on average appear to increase following strong levels of INS. Registration of thermal and nonlinear imaging fields within a couple of microns is key to enable direct correlation of temperature-time dynamics with cell functionality since cells are not readily visible with thermal imaging.

To practically extend this imaging platform, cell viability markers can easily be substituted for calcium sensors, voltage probes, FRET constructs, or gene transcription assays, alongside hyperspectral or bio-orthogonal SRS approaches, to systematically study the functional impact of INS on other aspects of cellular physiology. SRS often provides relatively nonspecific biomolecular information relative to fluorescence imaging strategies; combining functional fluorescence and SRS imaging strategies during INS can expand the current understanding of the impact of cellular physiological dynamics on SRS signals. Similar approaches can be used to understand the effects of rapid temperature changes on more specific aspects of cellular physiology to improve our general understanding of the impact of thermodynamics on cell physiology.

Beyond INS, similar approaches as demonstrated in Fig. 6 may readily be adapted to study laser tissue interactions associated with optogenetic or nanoplasmonic neuromodulation, laser preconditioning of immunological response, and photodynamic therapy of infectious or cancerous model systems where temperature changes may have a substantial impact. These types of experiments are readily enabled only by an imaging platform that integrates nonlinear and thermal microscopy with high enough spatial and temporal resolution capabilities to study the processes at hand. While neuroscientific questions provide a valuable benchmark in terms of evaluating imaging speed in the context of biology, this platform is just as applicable to other areas of biomedical research.

Undoubtedly, an imaging platform that simultaneously integrates thermal and nonlinear imaging would be practically important in answering many questions in the afore mentioned disciplines. It becomes technically difficult to accommodate the disparity in optical detection wavelengths for use in a broad range of cellular, tissue, and live animal preparations. Tradeoffs between sample flexibility and optical access were major considerations in the imaging system’s three-armed design. Inherent hardware limitations prevent the design a truly simultaneous multimodal nonlinear and thermal imaging system. Nonetheless, the MANTIS platform offers a novel approach to creatively answer fundamental and translationally relevant questions about biology and thermodynamics which are applicable broadly to biomedical research.

## Conclusion

We present a novel and flexible multimodal optical imaging system that combines nonlinear and thermal microscopies for biomedical imaging applications. Our platform can be used to overlay functional, structural, and biochemical information from a single specimen tracked over time at high speed and spatial resolution. Such approaches can be applied to study dynamic biophysical processes in cells and tissues, in vitro and in vivo. In conjunction with continually expanding molecular biology tools, this instrument and related imaging methods will aid in studying physiological and biochemical processes from multiple perspectives in many fields including neuroscience, cancer biology, metabolic disease, tissue biomechanics, and much more.

## Funding

Funding for this work was provided by the following grants: AFOSR DURIP FA9550-15-1-0328, AFOSR FA9550-14-1-0303, AFOSR FA 9550-17-1-0374. WRA was supported through the ASEE NDSEG Fellowship.

## Acknowledgements

The authors would also like to thank Dr. Bryan Millis, Dr. Christine O’Brien, and Dr. Andrea Locke for their feedback and guidance on the manuscript. The authors would also like to thank Dr. Matthew Bersi, Joshua Bender, and Dr. W David Merryman for providing the porcine mitral valve samples.

## Author Contributions

AMJ, EDJ, and JB conceived the idea for the manuscript. WRA, BM, EL, SP, JB and AMJ designed, and built the imaging system. AMJ and EDJ secured funding support for the published work. MW performed *in vivo* surgical preparations and assisted in imaging. RG, GT, JLJ, and JBF provided samples and experimental guidance imaging tissue samples. WRA assisted in all sample preparations, performed all imaging experiments, image processing, data analysis, and prepared the manuscript. All authors contributed to editing manuscript.

## Data Access

Any raw data and processing scripts are readily available from the corresponding authors upon request.

## Conflicts of Interest Statement

BM, EL, SP, and JB are employees of Thorlabs Imaging Research, LLC. The remaining authors report no additional conflicts of interest.

## Supplemental Materials

**Table S1:**
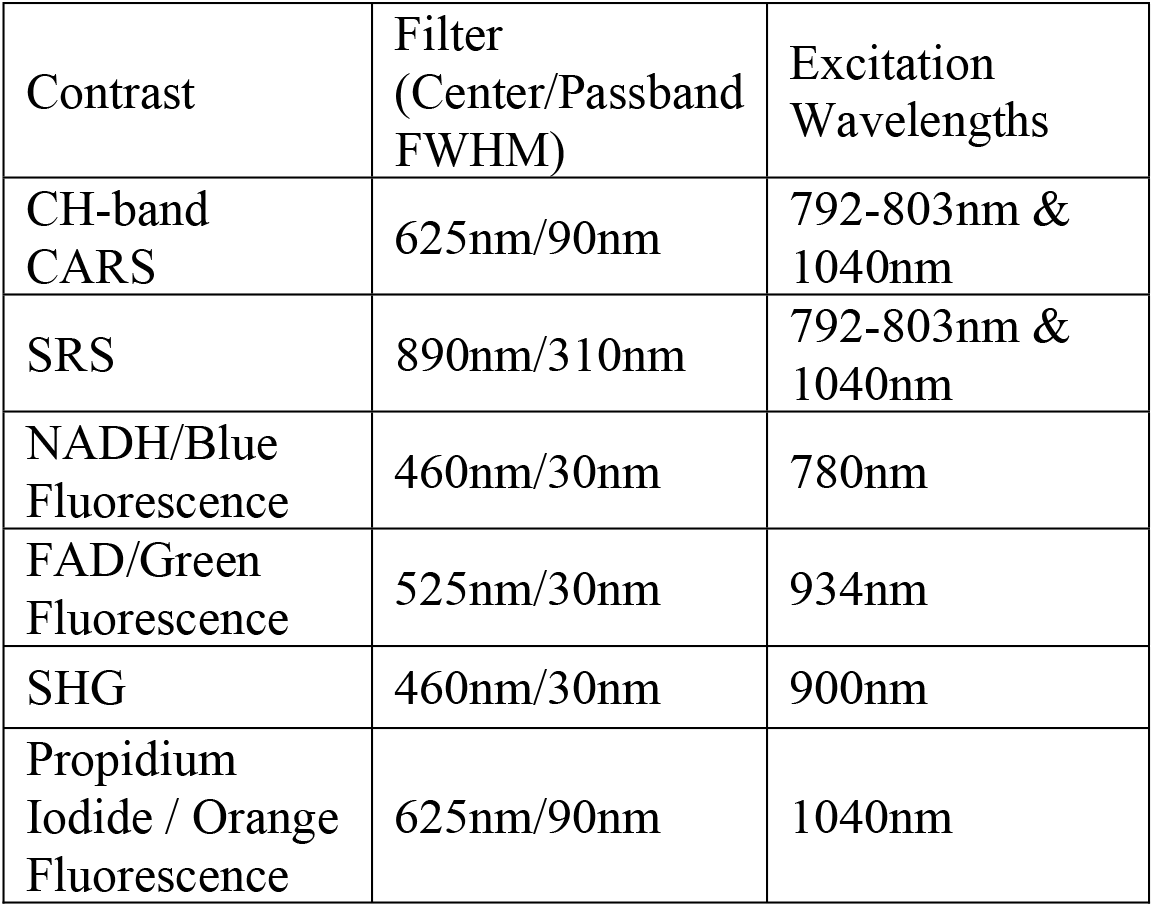
Summary of emission filters and excitation wavelengths used for multimodal imaging. All filters obtained from Semrock (Brattleboro, VT, USA).

**Table S2:** Summary of measured system resolutions across multiple imaging modalities.

| Modality | Lateral<br>Resolution<br>($\mu\text{m}$) | Axial<br>Resolution<br>($\mu\text{m}$) |
| --- | --- | --- |
| CARS | 0.632 | 3.009 |
| SRS | 0.833 | 3.136 |
| MPF | 0.359 | 1.502 |
| SHG | 0.388 | X |
| Thermal | 6.9 | X |

**Table S3:**
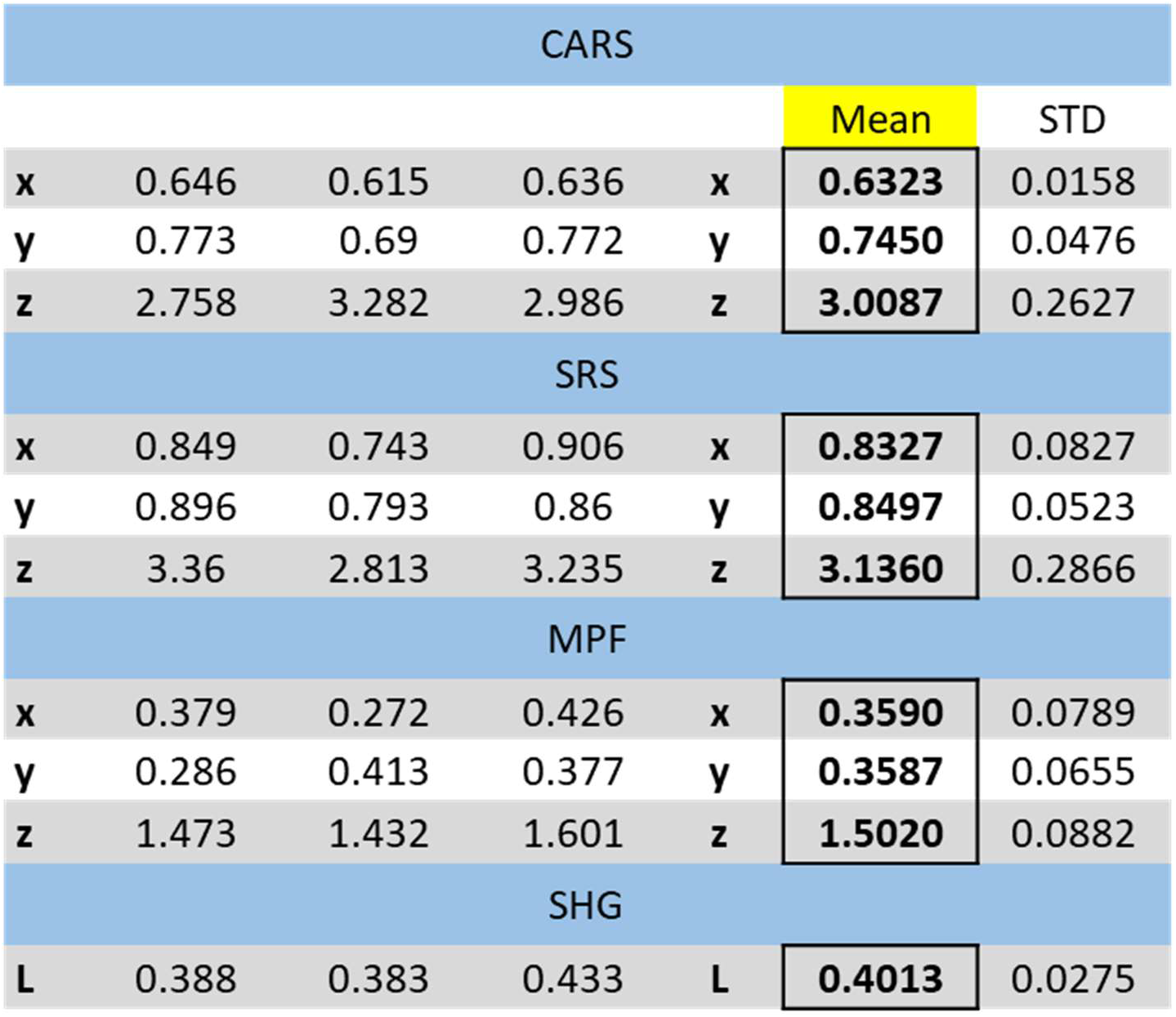
Detailed summary of point spread function calculations for nonlinear imaging modalities.

**Figure S1:**
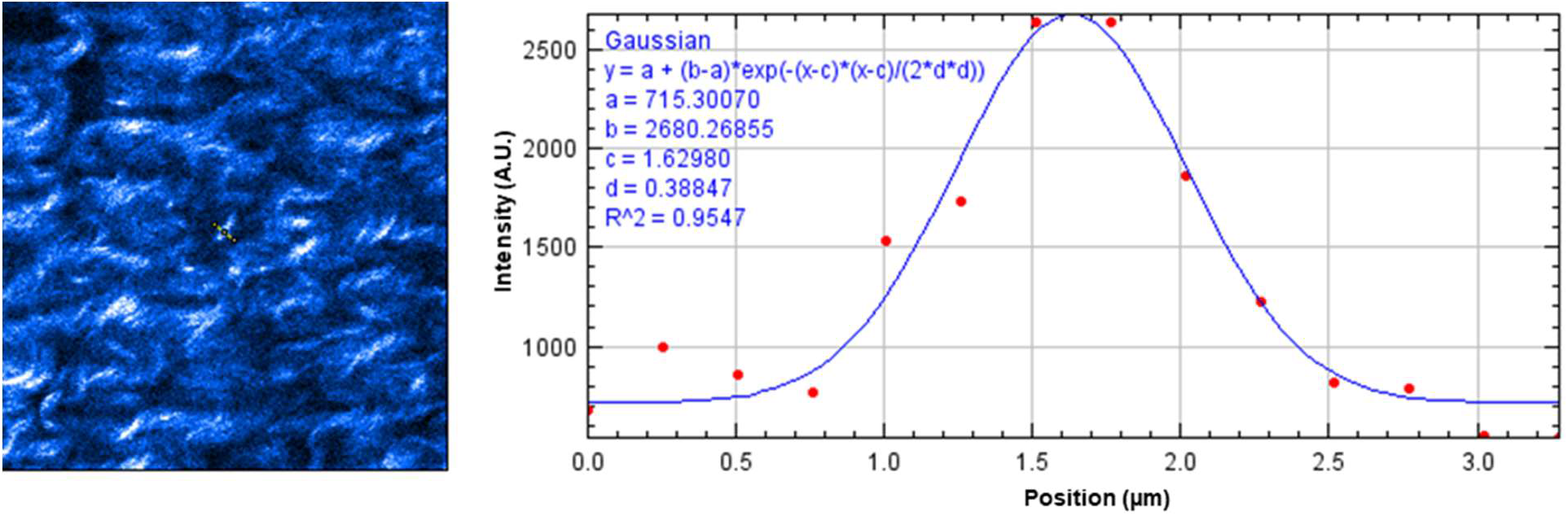
Sample Lateral Resolution Characterization for SHG imaging in a porcine mitral valve sample. Gaussian fitting performed in FIJI.

**Figure S2:**
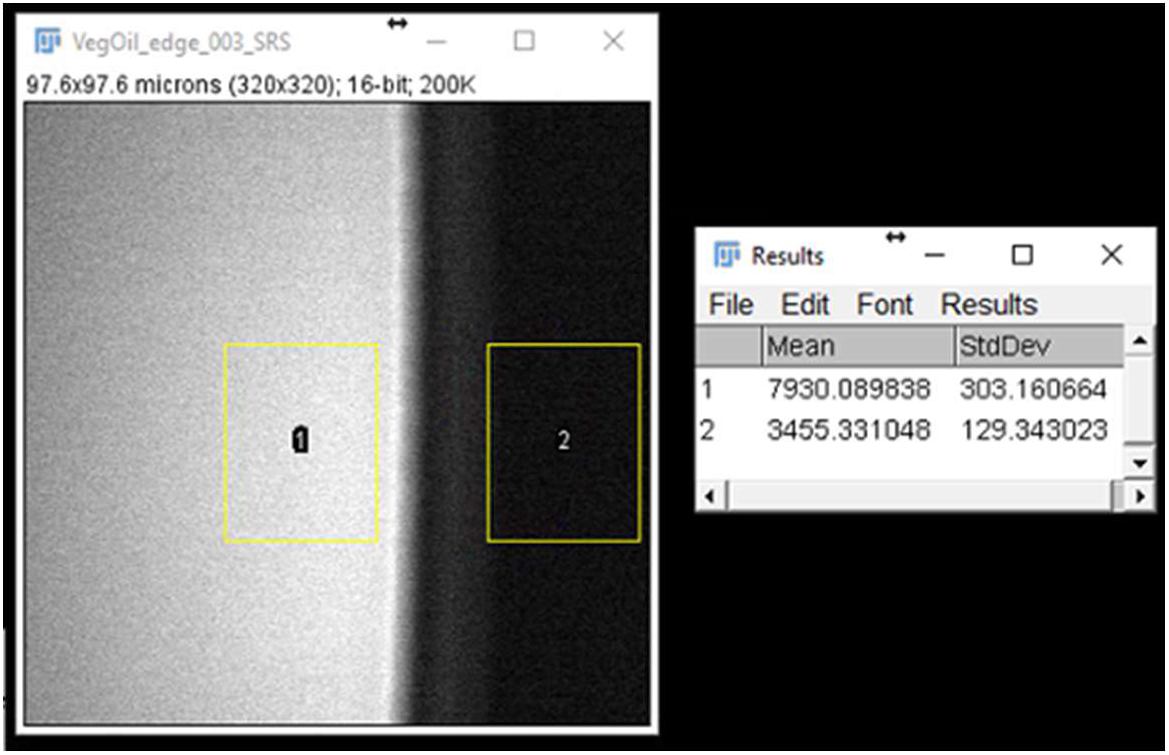
Signal to Noise calculation with a Vegetable Oil meniscus in FIJI. SNR was calculated to be 34.6.

**Figure S3:**
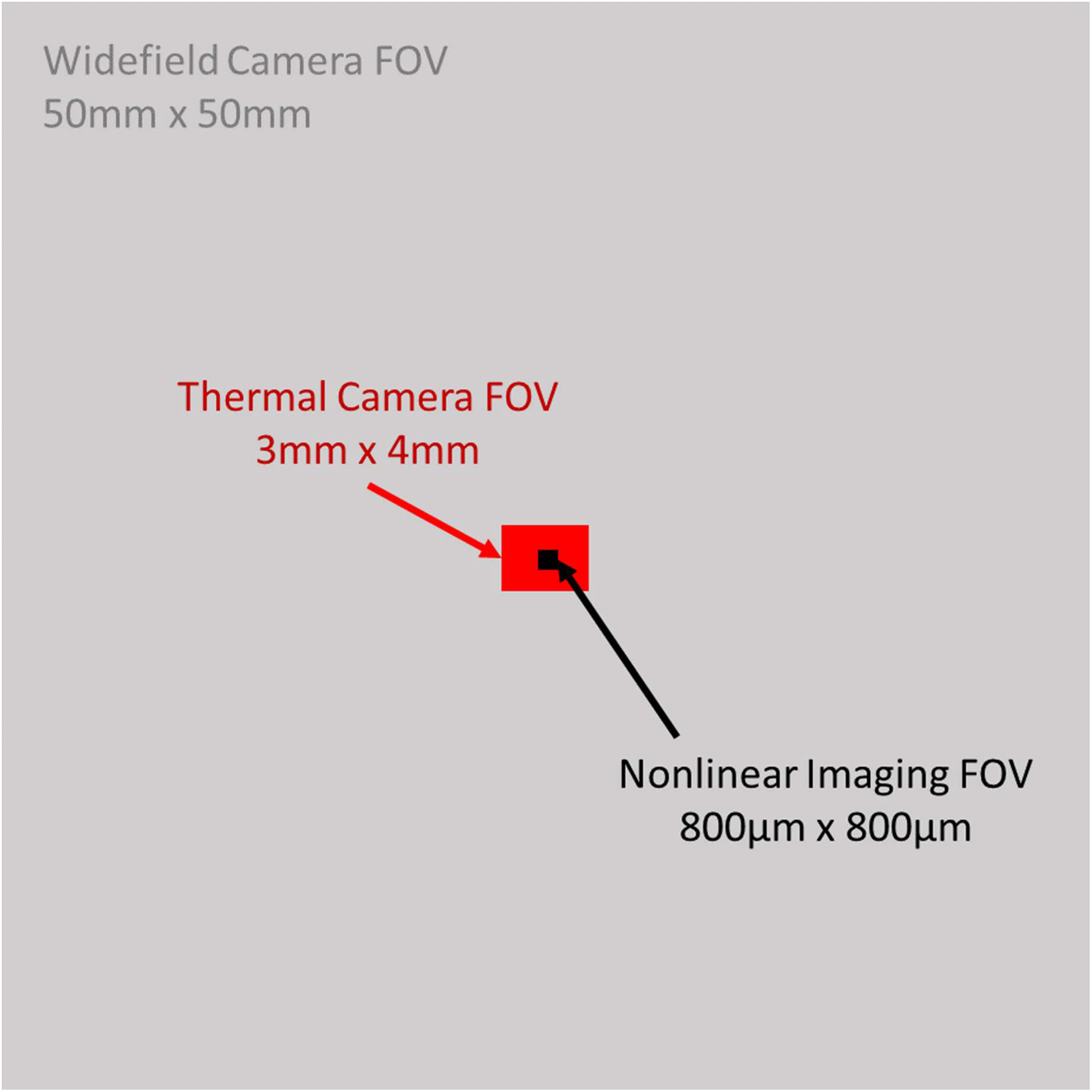
To-scale representation of the overlap of multimodal imaging fields of view for the 3 imaging arms of MANTIS.

**Figure S4:**
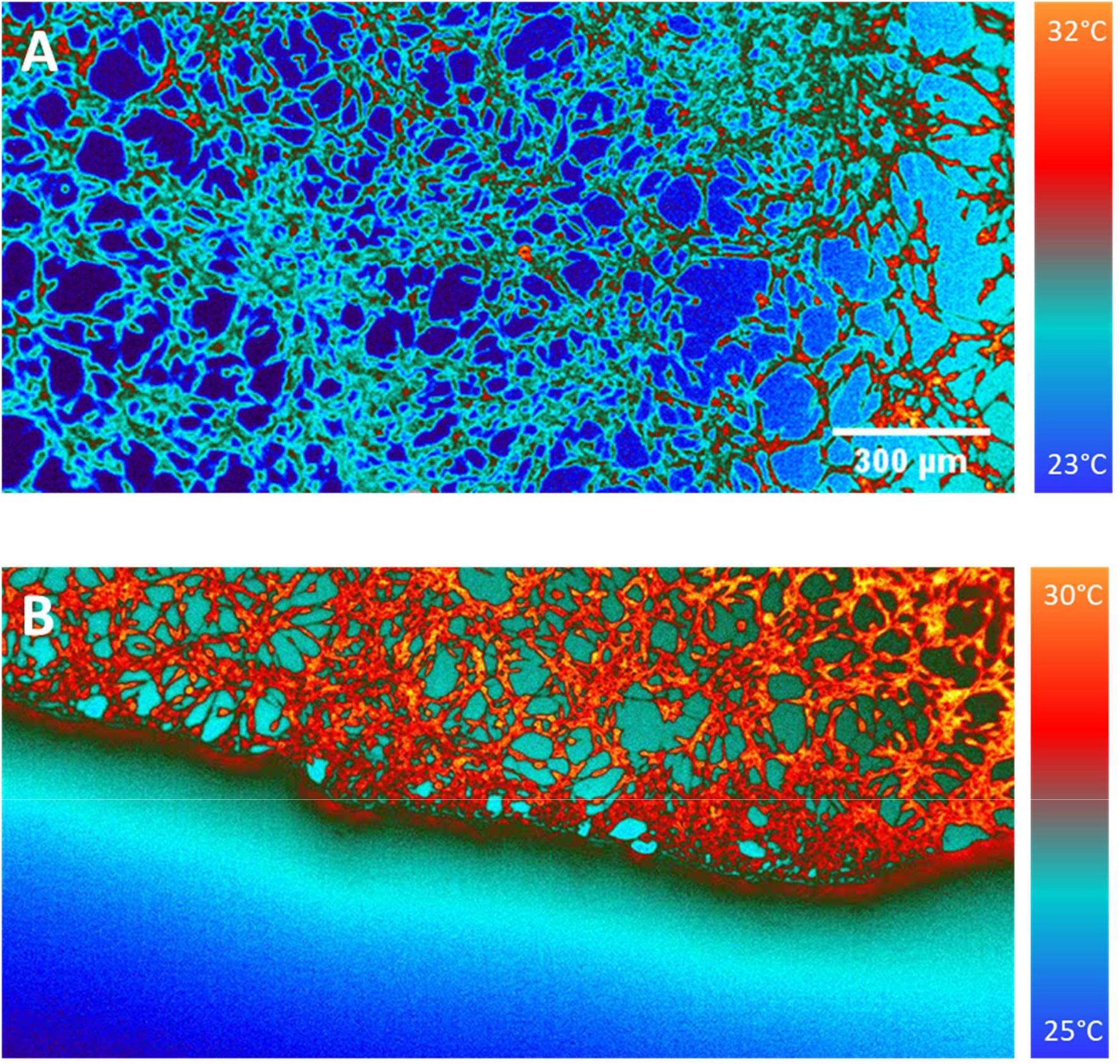
Thermal images of cultured 3T3 Fibroblasts. A) Cells imaged without any aqueous medium. B) Image of aqueous medium front advancing over cells. The absorption of water in the short-wave infrared is high, making cell culture medium difficult to image through with blackbody thermal contrast.

**Figure S5:**
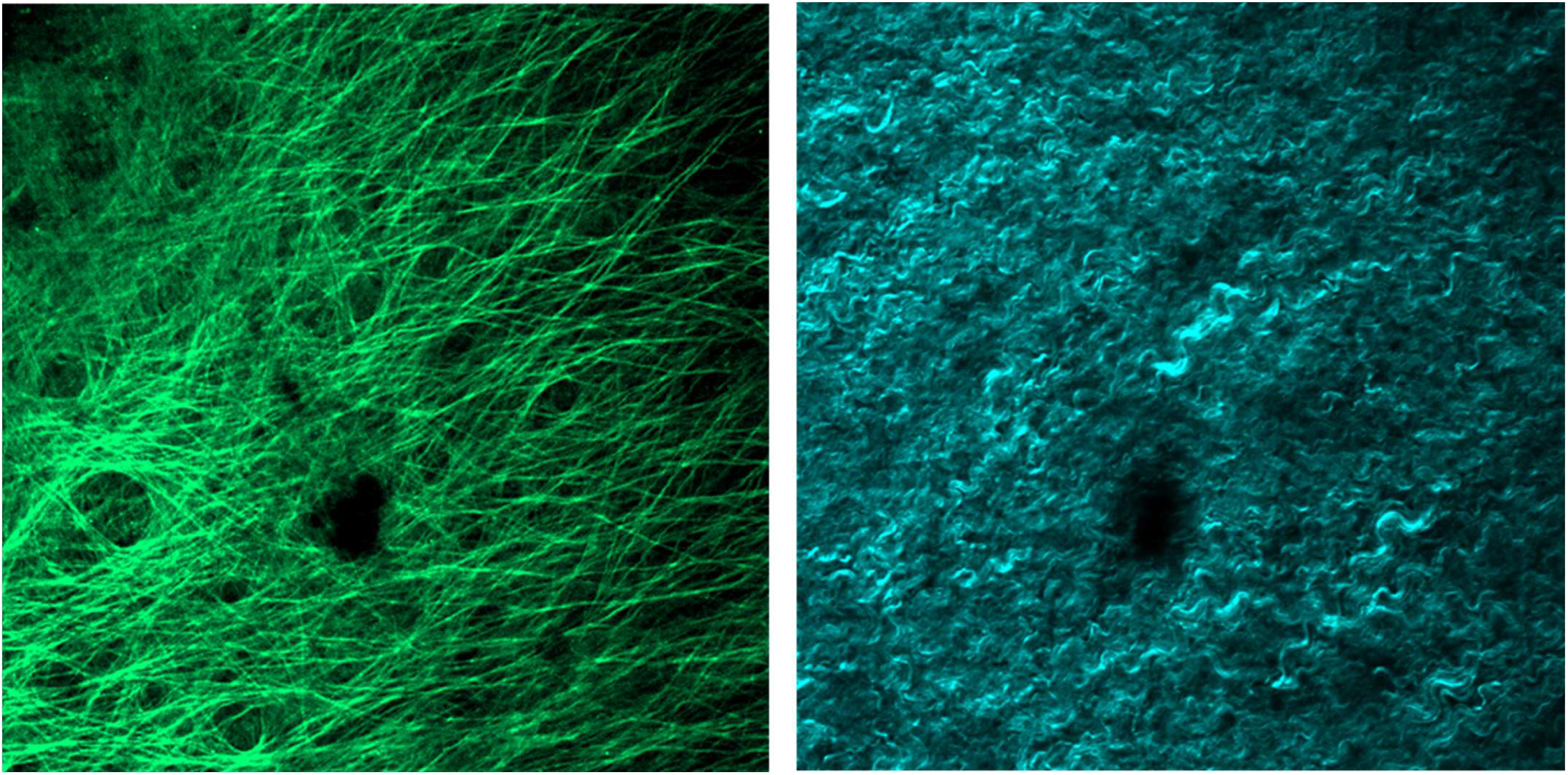
Porcine Mitral Valve imaging with endogenous autofluorescence from elastin (left) and collagen SHG (right). Image width is 520-μm.

**Figure S6:**
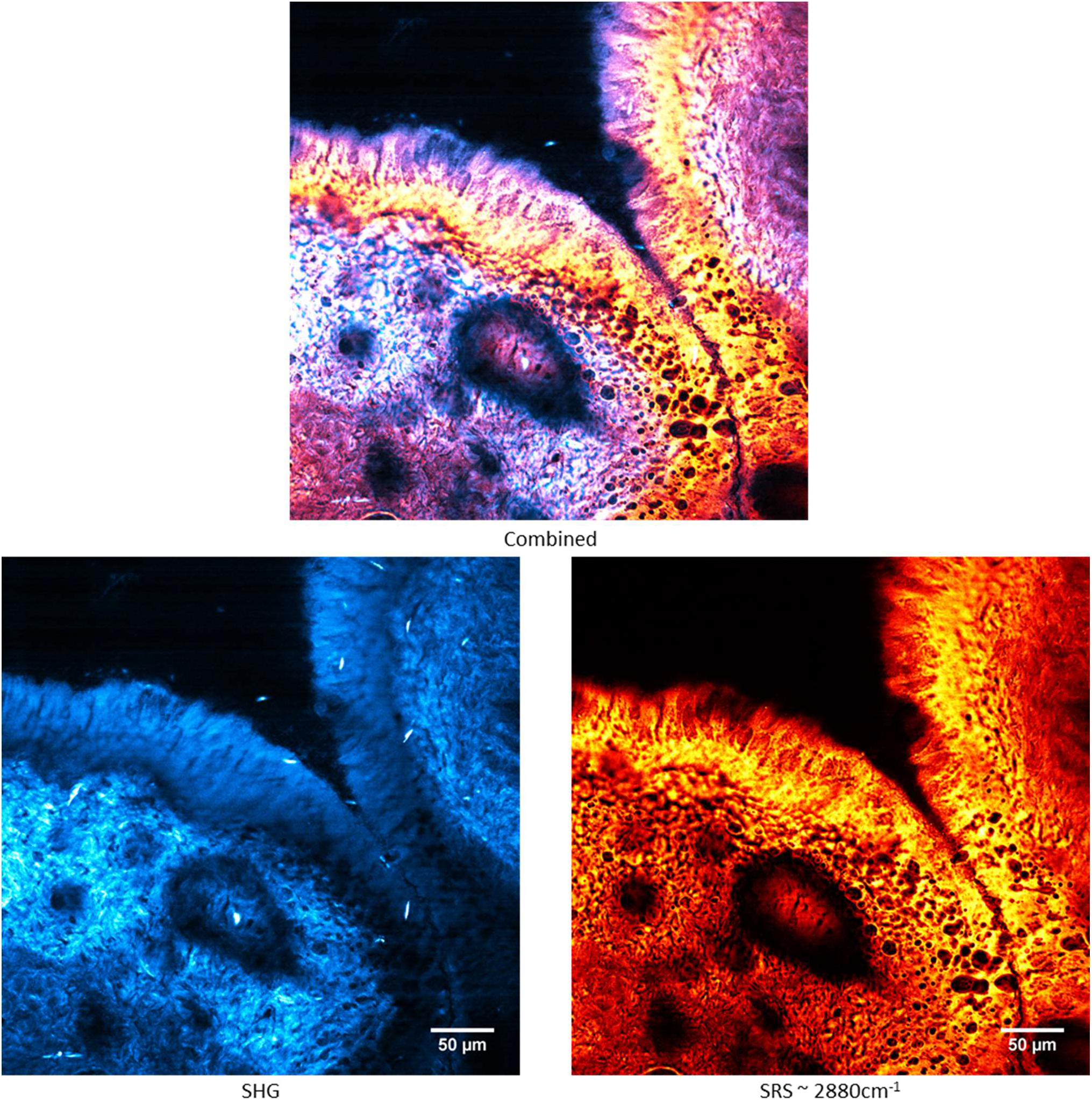
Simultaneous SHG (cyan) and SRS (2880-cm-1, lipid dominant resonance, orange) imaging of an unstained murine cervix unfixed frozen section. The stroma (dense in collagen) and epithelium are easily discernable based on relative concentrations on ratio of SHG and SRS signals. Blood vessels lamina propria are also visible with SHG contrast.

**Figure S7:**
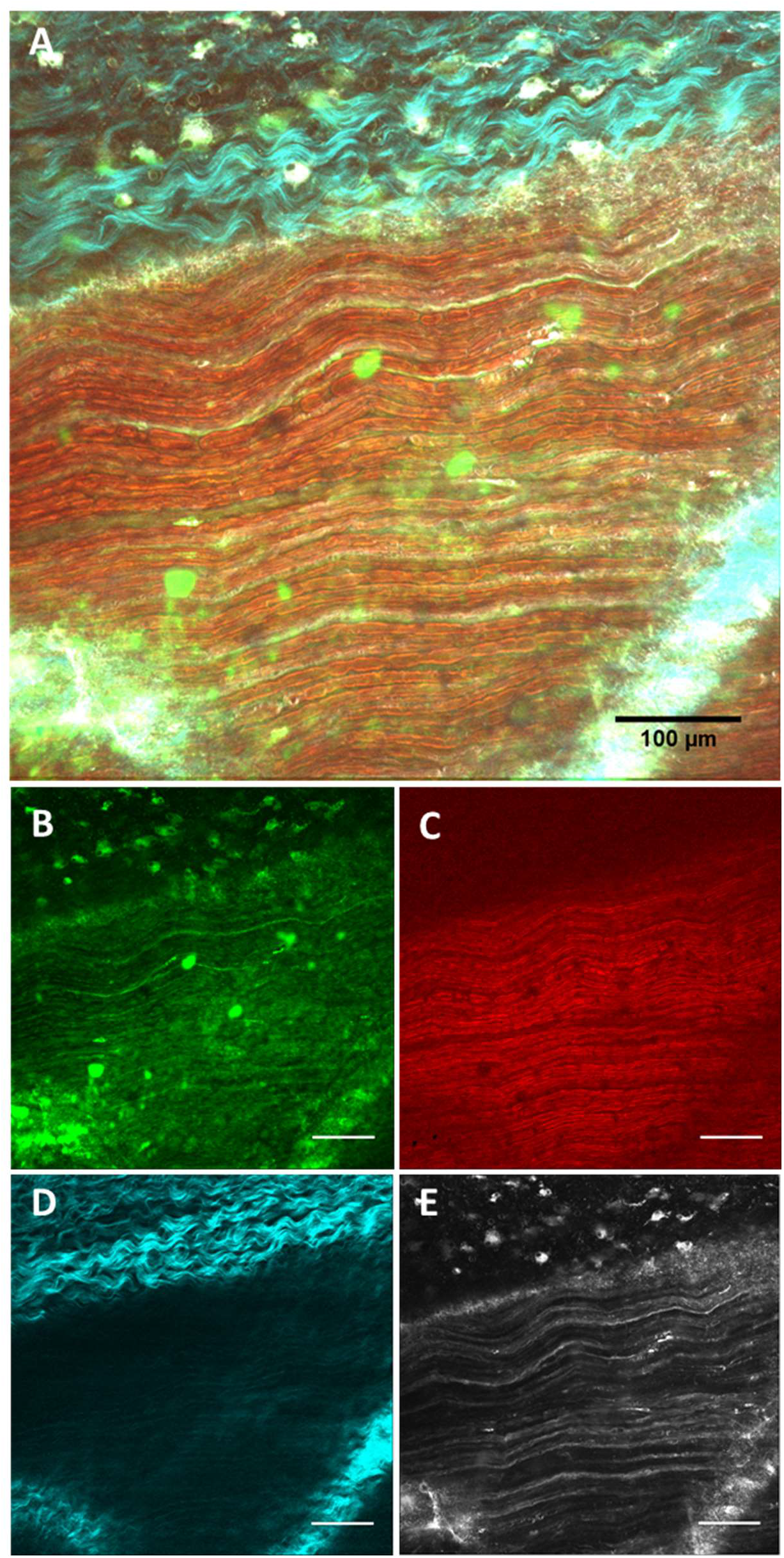
Composite multimode images of an *ex vivo* rat sciatic nerve. (A) CARS signal at 2927cm^−1^ (B, green), SRS signal at 2927cm^−1^ (C, red), SHG signal (D, cyan), and multiphoton fluorescence of FluoroMyelin Green (E, Grey). All scale bars are 100um.

**Figure S8:**
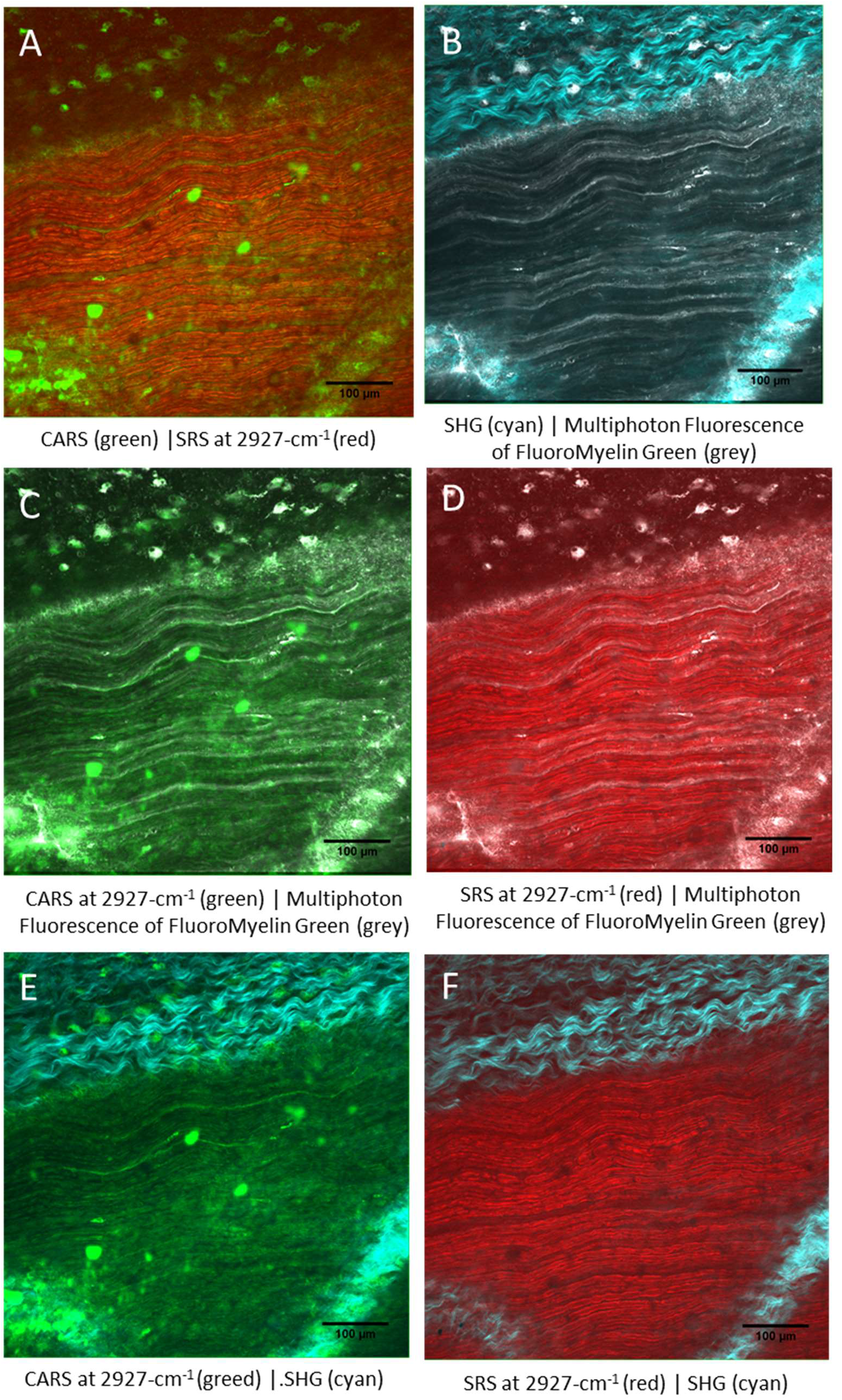
Ex vivo rat sciatic nerve samples imaged with multimodal nonlinear imaging from Supp. Fig. S7. Two different modalities can be combined in different ways to visualize tissue structure.

**Figure S9:**
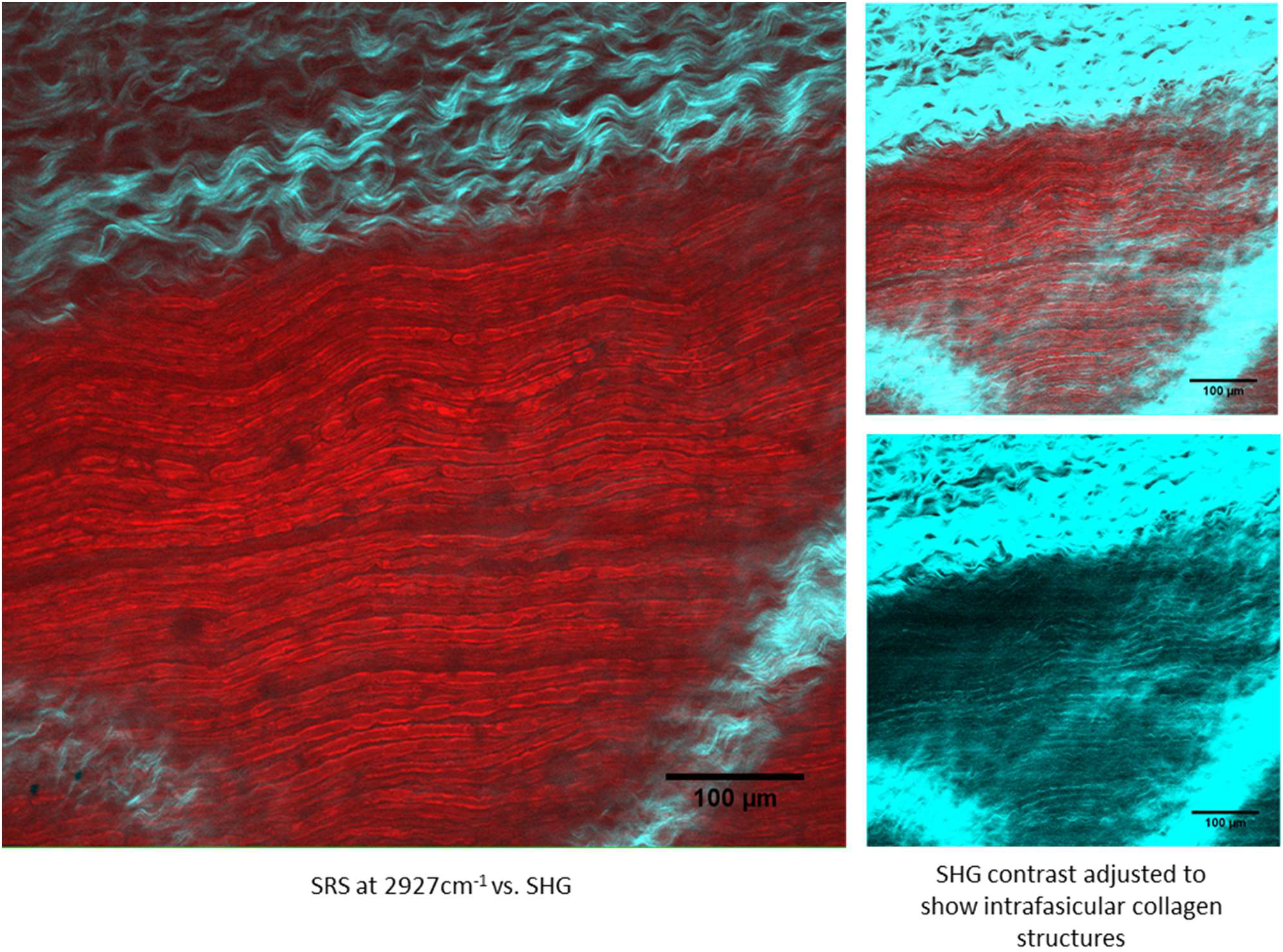
Supplementary Figure 9: Ex vivo rat sciatic nerve samples imaged with SRS (red, myelin at 2927cm-1) and SHG (cyan, collagen). Rescaled SHG images are shown to highlight intrafasicular collagen, which is in lower abundance and overall signal then epineurial collagen.

## Notes

### Summary of Updates

The manuscript file is being updated following peer review.

